# Nuclei are mobile processors enabling specialization in a gigantic single-celled syncytium

**DOI:** 10.1101/2021.04.29.441915

**Authors:** Tobias Gerber, Cristina Loureiro, Nico Schramma, Siyu Chen, Akanksha Jain, Anne Weber, Anne Weigert, Malgorzata Santel, Karen Alim, Barbara Treutlein, J. Gray Camp

## Abstract

In multicellular organisms, the specification, coordination, and compartmentalization of cell types enable the formation of complex body plans. However, some eukaryotic protists such as slime molds generate diverse and complex structures while remaining in a multinucleated syncytial state. It is unknown if different regions of these giant syncytial cells have distinct transcriptional responses to environmental encounters, and if nuclei within the cell diversify into heterogeneous states. Here we performed spatial transcriptome analysis of the slime mold *Physarum polycephalum* in the plasmodium state under different environmental conditions, and used single-nucleus RNA-sequencing to dissect gene expression heterogeneity among nuclei. Our data identifies transcriptome regionality in the organism that associates with proliferation, syncytial substructures, and localized environmental conditions. Further, we find that nuclei are heterogenous in their transcriptional profile, and may process local signals within the plasmodium to coordinate cell growth, metabolism, and reproduction. To understand how nuclei variation within the syncytium compares to heterogeneity in single-nucleated cells, we analyzed states in single *Physarum* amoebal cells. We observed amoebal cell states at different stages of mitosis and meiosis, and identified cytokinetic features that are specific to nuclei divisions within the syncytium. Notably, we do not find evidence for predefined transcriptomic states in the amoebae that are observed in the syncytium. Our data shows that a single-celled slime mold can control its gene expression in a region-specific manner while lacking cellular compartmentalization, and suggests that nuclei are mobile processors facilitating local specialized functions. More broadly, slime molds offer the extraordinary opportunity to explore how organisms can evolve regulatory mechanisms to divide labor, specialize, balance competition with cooperation, and perform other foundational principles that govern the logic of life.

## Introduction

Animals and other multicellular organisms are composed of diverse, compartmentalized cell types that perform specialized functions. Coordinated behavior between cells has led to the evolution of complex body plans with specialized organs. Furthermore, intercellular interactions between cell types are required to maintain a balanced physiology in a dynamic environment. The need for coordinated behaviour over large spatial domains led in some cases to the formation of multinucleated cells, also called syncytia. Examples are skeletal muscles of animals (reviewed in (1)), the developing endosperm in many plants (coenocytes) (2), multinucleated hyphae in different fungi (3), or the plasmodia of many protists (4–7). The syncytium, a cytoplasmic mass containing numerous nuclei that store genetic material including deoxyribonucleic acid (DNA), originates either through merging of cells or through nuclei divisions lacking an accompanying cell division. An example for the latter mode of syncytia formation are the plasmodia of many slime molds. Interestingly, such acellular slime molds are able to generate complex and dynamic body structures covering tens of centimeters or more while lacking compartmentalization of nuclei into discrete cellular units. Such slime molds show that multicellularity is not necessary to form specialized structures with specific functions. The underlying molecular processes that lead to syncytium formation and enable different subdomains to respond to dynamic environments are poorly understood.

The slime mold *Physarum polycephalum* has a multi-phasic life cycle where uninucleated haploid amoeba merge to form a diploid zygote which grows into a gigantic protoplasmic syncytium, called a plasmodium (Fig. 1A) (8). The plasmodium forms a network-like structure to span and connect food sources and other amenities (9) (Fig. 1B). The morphologically complex *Physarum* plasmodium can grow meters in area, exhibit diverse phenotypic behaviors, and dynamically respond to environmental conditions that it encounters (10). The plasmodium’s network-like body plan consists of interlaced tubes of varying diameters, and it was recently shown that these tubes grow and shrink in diameter in response to a nutrient source, thereby functioning to imprint the nutrient’s location in the tube diameter hierarchy (11). The syncytium is composed of tens to millions of nuclei depending on the plasmodium size, and the nuclei are thought to divide synchronously as it grows (12, 13). Studies have described that nuclei can be mobile and flow dynamically within cytoplasmic spaces, moving throughout the plasmodium (14, 15) with up to 1.3 mm per second which is among the fastest cytoplasmic flows ever measured (16, 17). It is not known if nuclei diversify into heterogeneous states, if region-specific gene expression modules are established that guide the formation of the complex morphological structures, and if modules enable local response to environmental stimuli.

**Fig. 1.**
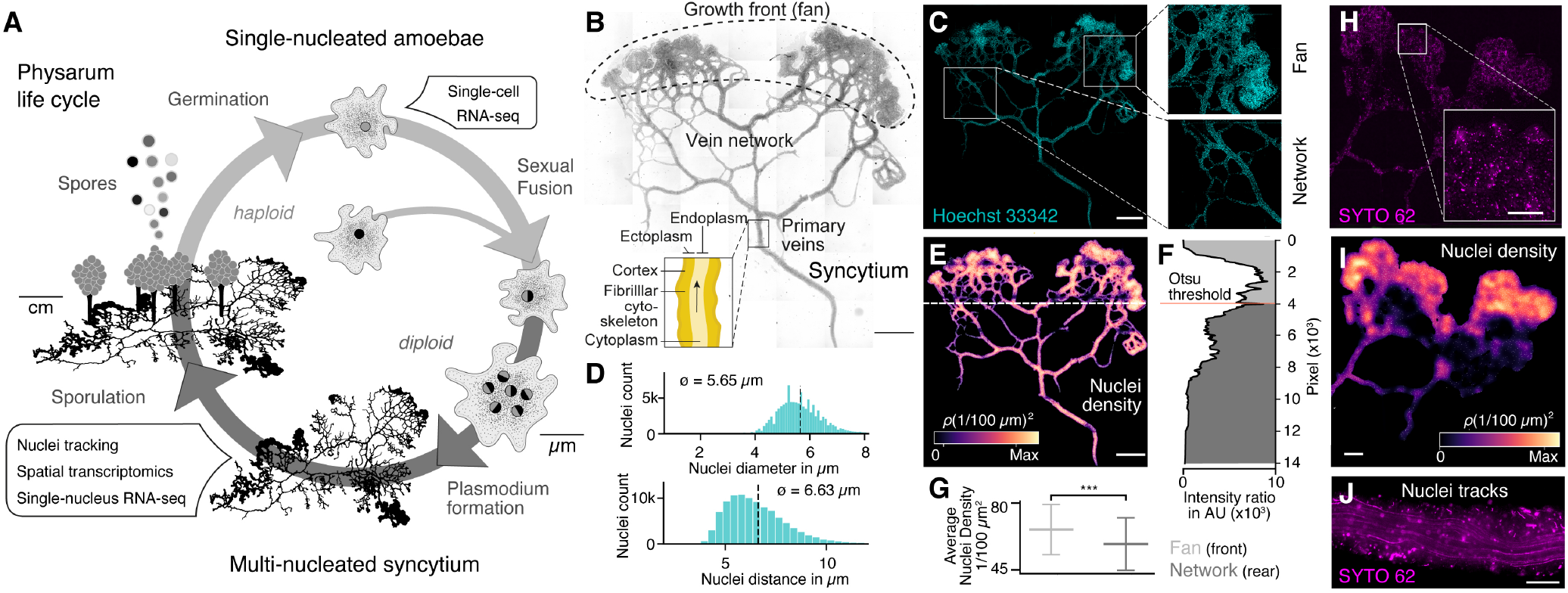
The slime mold *Physarum polycephalum* forms a multinucleated syncytium with mobile nuclei. (**A**) Simplified life cycle of *Physarum polycephalum* indicating which experiments were performed in this study. (**B**) Plasmodium of *Physarum polycephalum* grows as a hierarchical network composed of contractile veins and a microstructured growth front. Tiled brightfield image of a fixed plasmodium is shown as reference. Scale bar: 1000 μm. (**C**) Hoechst stained nuclei across the syncytium from (B), a growth front (top right) and a primary vein (bottom right). Scale bar: 1000 μm. (**D**) Histograms of nuclei diameter (top) and density (bottom) estimates are shown. Mean values are provided as insets. (**E**) Density map of nuclei after StarDist segmentation and kernel density estimation shows inhomogeneously distributed nuclei (see Methods for further details). Scale bar: 1000 μm. (**F**) Averaged relative density estimates across the x-axis of the plasmodium are visualized along the y-axis of the plasmodium. A threshold by Otsu (19) was estimated to unbiasedly find a change in the density distribution separating the plasmodium in a network and a fan region. (**G**) Boxplots visualize the average nuclei density for the fan or network region identified in (F). (**H**) Tiled life image of a plasmodium stained with SYTO62 and a magnified fan region (inset). Scale bar: 500 μm. (**I**) Density map of nuclei shows inhomogeneously distributed nuclei. Scale bar: 500 μm. (**J**) Maximum Intensity projection over a period of 5 seconds of nuclei motion visualized by streamlines. See suppl. Video 1. Scale bar: 100 μm.

To address these outstanding questions, we performed spatially-resolved RNA-seq and single-nucleus RNA-seq on multiple slime mold plasmodia under different growth conditions. The data enabled mapping of gene expression profiles onto different structures within the plasmodium and exploration of localized transcriptome responses which correlated with nuclei heterogeneity. Taken together, our data suggests that nuclei within *Physarum* are mobile and can integrate local signals to coordinate a transcriptional response to dynamic environmental conditions, which enables the syncytium to locally change behavior and morphology.

## Results

### Nuclei are distributed throughout the slime mold syncytium and can be mobile

We pursued several imaging experiments on fixed and live plasmodial cultures to better understand nuclei size, location, and dynamics within different substructures of the syncytium (Fig. 1B and Fig. S1). We first grew plasmodial cultures on phytagel plates, fixed the plasmodium with paraformaldehyde, and stained nuclei with Hoechst 33342. We observed tens of thousands of nuclei distributed in the growth front (fan) and network veins (Fig. 1C). We found that the nuclei have an estimated mean size of 5.65 μm and a mean inter-nuclei distance of 6.63 μm (Fig. 1D). In particular, the maximum distance between two nearest nuclei is 33.9 μm, but 99.9% of the nuclei are within a distance of 12 μm apart from each other. We used a convolutional neural-network to segment nuclei (18), which revealed a heterogeneous distribution of nuclei across the plasmodium (Fig. 1E). The mass differences across the plasmodium were used to unbiasedly separate the plasmodium into a fan and a network part by estimating the Otsu threshold (Fig. 1F) (19) and we observed that the growing fronts show significantly increased nuclei densities compared to network veins (Fig. 1G).

To explore nuclei dynamics in the syncytium, we stained live plasmodia using a fluorescent d ye (SYTO 6 2) and imaged vein networks and fan regions in the syncytium over 100 seconds (Fig. 1H). Nuclei segmentation (18) confirmed a heterogeneous distribution of nuclei, with an increased nuclei-density within the growth front of the slime mold similar to what was observed in the fixed plasmodium (Fig. 1I). Within the same imaging field, we observed immobile and mobile nuclei in close proximity. Mobile nuclei are advected by the local shuttle flow in veins of the network (Fig. 1J, suppl. Video 1) and the foraging front (see suppl. Video 2). Strikingly, advected and immobile nuclei coexist in the syncytium and can get trapped in the ectoplasm or be released into the peristaltic flow of the endoplasm. Altogether, these data show that nuclei are distributed throughout the plasmodial syncytium, and that they are maneuvered as structures form and grow.

### Spatial RNA-seq uncovers morphology- and region-specific gene expression profiles in the slime mold syncytium

To determine if there is transcriptional heterogeneity in different domains and structures within the slime mold, we performed spatial RNA-seq measurements across individual slime mold plasmodia (SM1-4, Fig. 2A and Fig. S2A; see Fig. S3 for additional experiments). All slime molds (SM1-4) originated from the same micro-plasmodium culture and were washed with nutrient-free medium to ensure similar starting conditions prior to growth on standard 384-well plate covers coated with a thin agar layer. SM1 and SM2 were collected after 20 hours of growth whereas for SM3 and SM4, 1 oat flake was placed 2 cm away from the initiating plasmodium. After an additional 3 hours of culture, growth fronts in both SM3 and SM4 were in close proximity to the oat flake (Fig. S2A). At this point, SM3 was collected for spatial RNA-seq, whereas SM4 was allowed to cover and assimilate the oat, and grow for an additional 4 hours prior to collection for spatial RNA-seq (Fig. S2A). Spatially restricted plasmodium sampling was achieved by placing slime molds onto a 384-well plate reservoir followed by centrifugation such that position information was preserved in 4 millimeter square grids (Fig. S2B). Bulk transcriptome libraries were generated using the SMART-SEQ2 protocol (20) with small modifications (see Methods), barcoded and sequenced, and expression profiles were analyzed and mapped back to grids overlaying the original plasmodial structure. Spatially registered data from all slime molds were combined and a non-linear dimensionality reduction by uniform manifold approximation projection (UMAP) was performed (Fig. 2B). Grid clustering revealed distinct expression profiles from each slime mold (Fig. 2C). We identified differentially expressed (DE) genes between slime mold clusters, and found that SM3 grids showed the strongest difference in gene expression compared to the other plasmodia (Fig. S2C). This difference was driven by the expression of genes involved in signal transduction (DHKL) (21), chemotaxis (GEFA) (22), and cell shape polarity (TEA1) (23) (Fig. 2C). As noted above, a SM3 growth front was actively extending towards the oat flake at the moment of sampling (Fig. 2D) suggesting that the transcriptomic differences observed in this slime mold might originate from a chemotactic stimulus absent for the other slime molds. We note that SM2 also had elevated expression of other genes involved in chemo- and electrotaxis (GCA) (24) or responsiveness to a stimulus (Calpain-D/CALD), which is interesting as no food source or other stimulus was applied (Fig. 2C). In contrast, all grids of SM1 were enriched in cell cycle related genes (BUB1, MKI67, KIF3B) suggesting that SM1 was in the process of nuclei division (Fig. 2C; see also Fig. S3D)). These data suggest that transcriptome variation can be detected between slime molds grown from the same parent culture, but experiencing different conditions.

**Fig. 2.**
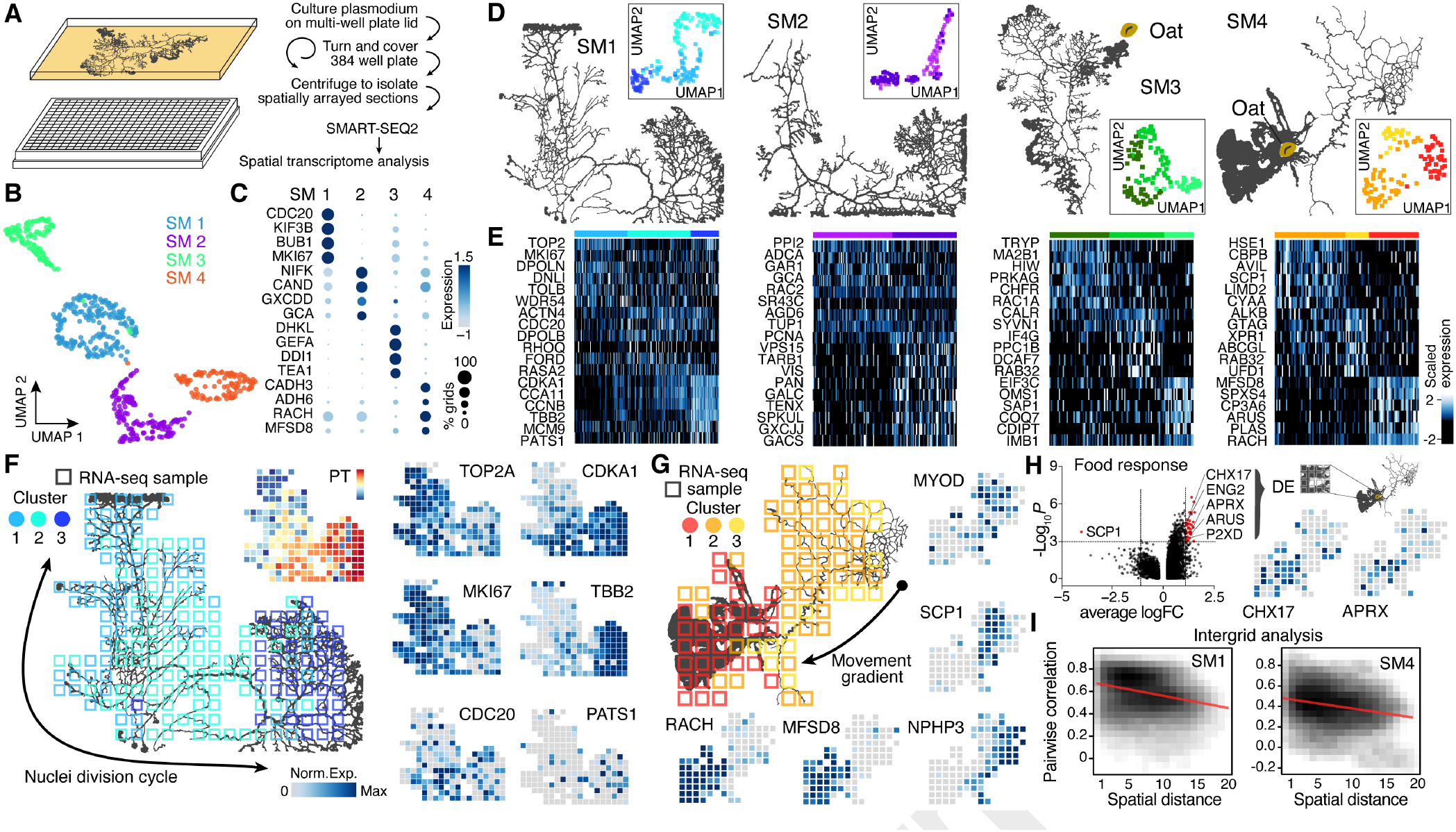
Spatial Transcriptomics reveals region-specific gene expression in the plasmodial slime mold syncytia. (**A**) Schematic of experimental set up. (**B**) Four slime mold plasmodia were grown overnight from a microplasmodia culture. UMAP embedding reveals general differences between slime molds. (**C**) Dotplot visualizes gene expression intensity (dot color) and frequency (dot size) for top slime mold specific marker genes, respectively. (**D**) Morphology overview of slime mold plasmodia investigated. SM1 and 2 were directly investigated after grown from microplasmodia culture whereas oat flakes were supplied to SM3 and 4 and experiments were performed either just before the slime mold reached the food (SM3) or 4 hours after it reached the oat flake and had started reexpanding (SM4). UMAP embeddings next to each plasmodium reveal gene expression differences across grids for individual slime molds. (**E**) Heatmap representations of gene expression intensities for cluster markers (rows) across all grids (columns). Grids are ordered by cluster identity shown in (D). (**F**) Sampling grid on the 384-well plate overlaid by the plasmodium of SM1 at the time when sampled (left). Different colors represent the clustering result from (D). Right: Marker genes from (E) are visualized as feature plots onto the sampling grid. (**G**) Sampling grid on the 384-well plate overlaid by the plasmodium of SM4 at the time when sampled (top left). Different colors represent the clustering result from (D). Bottom: Marker genes from (E) are visualized as feature plots onto the sampling grid. (**H**) Volcano plot (left) reveals DE genes specific to grids where the plasmodium was in direct contact with a food source (oat flake). Feature plots (right) visualize the spatial arrangement for 2 of the DE genes of the Volcano plot. (**I**) Density plots visualize the relation between the pairwise distance against the pairwise Pearson correlation across all samples of the grid of SM1 (left) and SM4 (right), respectively. Statistically significant negative correlations are identified and marked by a red line.

We next analyzed whether transcriptome heterogeneity also exists within individual slime mold syncytia. We performed dimensionality reduction for each individual slime mold spatial RNA-seq data, and visualized grid heterogeneity using UMAP (Fig. 2D). These analyses revealed substantial intra-syncytium heterogeneity, and we identified genes that were differentially expressed between clusters (Fig. 2E). Strikingly, we observed gene expression patterns that suggested coordinated intra-syncytial behaviors. For example, in SM1 a major source of heterogeneity was determined by differences in mitosis dynamics, as certain clusters expressed G2/M phase markers (TOP2A/MKI67), whereas other clusters expressed markers for other phases of the cell cycle. Mapping these transcriptomic profiles onto the respective grids of SM1 revealed a wave of molecular features evidencing nuclei division orchestration across the plasmodium (Fig. 2F; see also Fig. S2E-F), a phenomenon that was first observed in the beginning of the 20th century but has not been molecularly described (12, 13). Portions of the plasmodium show genes associated with G2M transitioning phase (TOP2A, MKI67) whereas the center of the slime mold is characterized by the anaphase promoting gene CDC20, and the bottom right portion by anaphase specific spindle and cytokinesis marker genes (CCNB, TBB2, AURAA) (Fig. 2F) suggesting that the wave of nuclei divisions initiated in the bottom right part of the slime mold (Cluster 3).

Similarly, when mapping heterogeneity observed in SM4 onto the spatial grids, we found that clusters distinguish morphological structures in the slime mold. For example, grid cluster 1 is associated with a growth front whereas other clusters overlap the network (Fig. 2G). We note that certain genes that generally distinguish SM4 from SM1-3, such as alcohol dehydrogenases ADH6 and CADH3, are ubiquitously expressed throughout this particular mold and may be involved in metabolizing oat compounds (Fig. 2C). Strikingly, there are genes enriched in the SM4 network region that associate with ectoplasm contraction (MYOD, SCP1), highlighting the role of the network in generating cytoplasmic flows across the plasmodium (Fig. 2G) (25). Interestingly, the sarcoplasmic calcium-binding protein 1 (SCP1) is also abundantly found in fast contracting muscles of invertebrates (26). The fan region, in contrast, is characterized by the gene racH which regulates actin filament polymerization in Dictyostelium (27). The expression of racH is linked with the coexpression of the major facilitator superfamily domain-containing protein 8 (MFSD8) which is a transmembrane carrier that transports solutes by using chemiosmotic ion gradients and plays a role in TORC1 signal transduction that controls the growth rate in yeast (28).

Another interesting aspect of SM4 is that it had recently assimilated an oat flake allowing us to test whether slime mold structures in direct contact with a nutrient source acquire a distinct gene expression profile. We selected a region of 4 × 4 grids covering the oat flake and performed a DE gene analysis against all other grids. Intriguingly, this analysis revealed genes with physiological links to nutrient uptake (Fig. 2H). For example, the putative K+/H+ exchanger CHX17 known to be expressed in Arabidopsis thaliana roots is involved in maintaining pH homeostasis (29). Similarly, the extracellular alkaline metalloprotease APRX (30) involved in casein degradation by pseudomonas and other psychrotolerant bacteria (31) is enriched in the region covering the oat flake.

Overall, for SM1, SM2 and SM4 there was a clear trend where the closer grids are in space, the more correlated they are in their transcriptome (Fig. 2I, Fig. S2E and G)). Altogether, this data demonstrates that the *Physarum* syncytium is able to generate localized gene expression patterns that are associated with structures, growth states, and environmental stimuli, and patterns are globally coordinated throughout the non-compartmentalized syncytia.

### Single-nucleus RNA-sequencing uncovers nuclei heterogeneity that correlates with spatial transcriptome dynamics and structure specialization

We next sought to determine if the spatial transcriptome differences observed in the syncytium are apparent at the level of single nuclei. We isolated nuclei from two plasmodia and performed single-nucleus RNA sequencing (snRNA-seq) using a droplet based single-cell genomics platform (10x Genomics) (Fig. 3A). The same plasmodium was sampled twice, first directly after an overnight plasmodium formation from a micro-plasmodium liquid culture (primary) and second after one week of culture with oat flakes (secondary, see Methods). Additionally, the 1-week secondary plasmodium was separated prior to nuclei isolation into a growth front-enriched (fan) and a network-enriched (network) sample (Fig. S4A). Syncytial nuclei are small with a diameter of around 5.6 μm (Fig. 1D), and we detected on average 591 transcripts (UMIs) per nucleus (Fig. S4B). Unsupervised nuclei transcriptome clustering and visualization of heterogeneity in an UMAP embedding revealed gradients of nuclei diversity rather than clearly distinct states in line with the lack of compartmentalization in the syncytium (Fig. 3B and E, (Fig. S4C-E). In the primary sample, we observed signatures suggesting a wave of nuclei division was in progress at the time of plasmodium sampling (Fig. 3B) and also noticed a nuclei cluster that specifically expressed racH, a gene we identified to be growth front-specific in the spatially resolved bulk data set (Fig. 2G). We extracted two clusters with strong mitosis signatures and ordered these nuclei along a cell cycle pseudotime (Fig. 3C and Fig. S4C). Comparison of this nuclei pseudotemporal trajectory with a pseudospatial ordering of grids from SM1, revealed strong concordance in wave-like mitotic gene expression signatures (Fig. 3D). This data supports that spatial gene expression heterogeneity can be observed at the level of single nuclei. The growth front and network samplings from the secondary plasmodium exhibited slightly more nuclei heterogeneity compared to the primary plasmodium (Fig. 3E and Fig. S4D). We observed unequal mixing of nuclei with certain clusters dominated by nuclei from either the growth front or the network region. Strikingly, the cluster dominated by growth front nuclei is marked by racH, whereas the network-dominated cluster is marked by SCP1 (Fig. 3F), in line with the spatial transcriptome data. Interestingly, there is a nuclei cluster which expresses a glutamate receptor (GRLE) and a Tyrosine-protein phosphatase (PTP2) both known to be involved in aggregation during fruiting body (sorocarp) development in the cellular slime mold Dictyostelium (32, 33) (Fig. 3F). We did not observe morphological signs of fruiting body formation, and analysis of previously identified sporulation-related genes (34) did not show clear enrichment in this cluster (Fig. S4E). These nuclei may be in a precursor state prior to induction of sporulation or sclerotium formation.

**Fig. 3.**
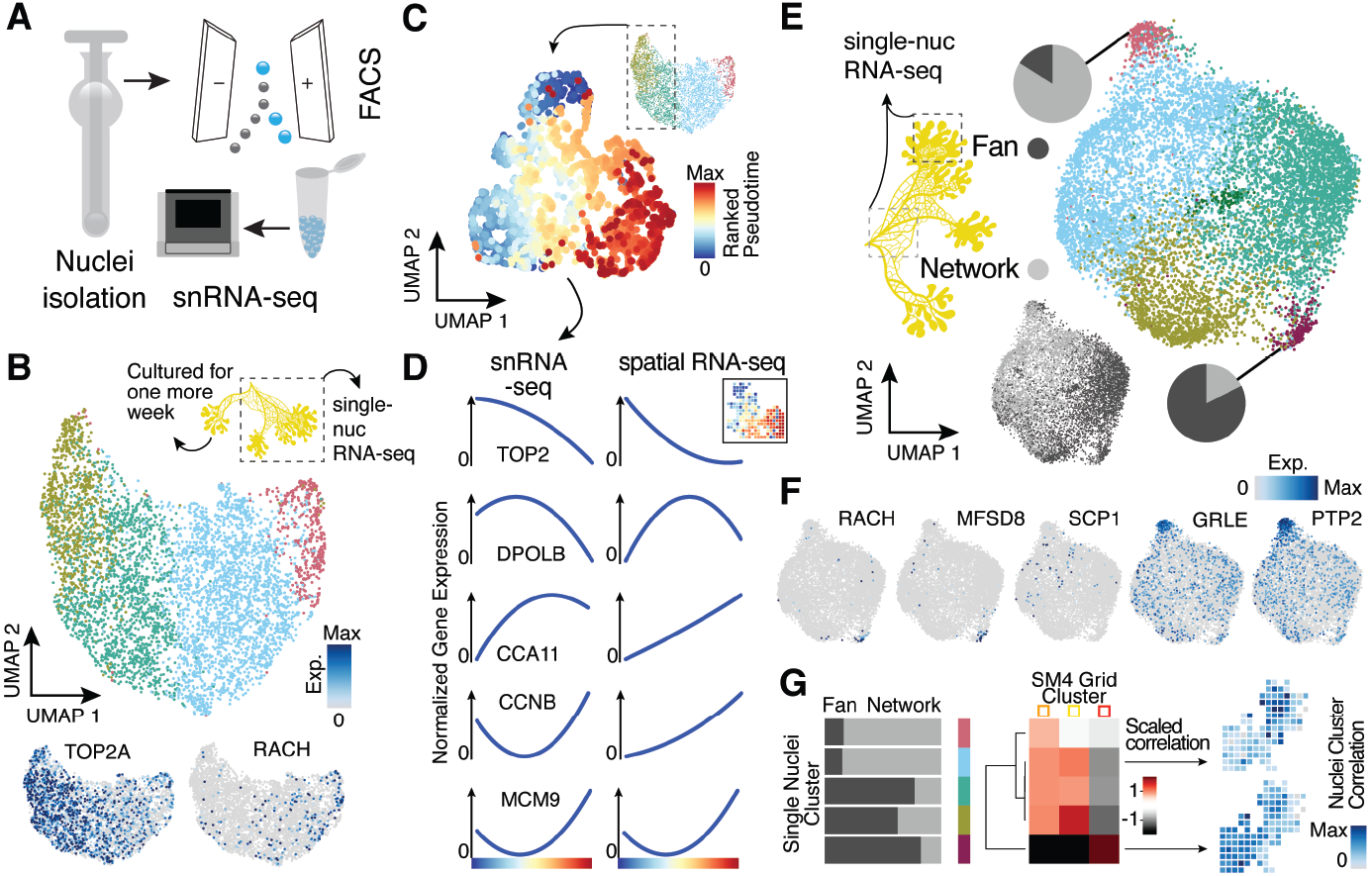
Single-nucleus RNA-sequencing uncovers nuclei diversification within the syncytia. (**A**) Nuclei extraction using a douncer with subsequent nuclei enrichment through FACS was performed prior to single-nucleus experiments. (**B**) UMAP embedding visualizes nuclei heterogeneity of a freshly grown plasmodium. DE marker genes are visualized as features on the embedding. (**C**) Pseudotemporal ordering of nuclei extracted from clusters with high TOP2A expression in (B). Extracted nuclei were re-embedded by UMAP with pseudotime ranks shown on the embedding. (**D**) Gene expression changes of differentially expressed genes across pseudotime are shown for snRNA-seq (left) and spatially resolved grid RNA-seq data (right). (**E**) UMAP embedding reveals nuclei heterogeneity of a 1-week old slime mold plasmodium with nuclei originating from different parts of the plasmodium. Pie charts reveal different nuclei proportions of 2 exemplary clusters. The UMAP inset encodes the origin of each nucleus by color. (**F**) Cluster marker genes for (E) are presented as feature plots on the embedding. (**G**) From left to right: Bar chart representing proportions of nuclei origin per cluster shown in (E). Heatmap representation of scaled correlation values between the pseudobulk gene expression per cluster of the plasmodium in (E) and the pseudobulk gene expression per cluster for SM4 (Fig. 2G). Arrows indicate for which clusters correlations against individual grids of SM4 were estimated with the result being presented as feature on the grid embedding.

We next determined if nuclei populations can be projected to spatial locations. We averaged expression values across single-nuclei clusters and across spatial transcriptome clusters from SM4 (Fig. 2G). We calculate correlations between nuclei and spatial clusters, and found that nuclei could be associated with different regions of the spatially segmented syncytium. Altogether, this data suggests that nuclei within the syncytium are transcriptionally heterogeneous, and that the particular transcriptome state associates with behaviour (e.g. proliferative growth) and location (growth front vs. network).

### Single-nucleated amoeba are heterogeneous and express genes distinct from nuclei within the single-celled syncytium

We next wanted to understand if the different nuclei states detected in the plasmodial syncytia are also observed in the single-nucleated *Physarum* amoeba stage and whether there are transcriptomic differences originating from the different nucleic division cycles (Fig. 4A-B). Towards this goal we characterized haploid *Physarum* amoeba cells using single-cell RNA-seq (Fig. 4A). We detected on average 14,525 transcripts per cell (Fig. 4C) and observed substantial variation in amoeba transcriptome states, identifying at least 6 distinct clusters at this resolution, with most of the heterogeneity being linked to cell cycle kinetics (35) (Fig. 4D). Interestingly, one cell cluster was marked by TOP6A, and we suspect that some haploid amoeba fuse and undergo meiosis (36, 37). These data illuminate amoeba cell states and reveal that substantial variation in cell states coexist within a *Physarum* single celled culture.

**Fig. 4.**
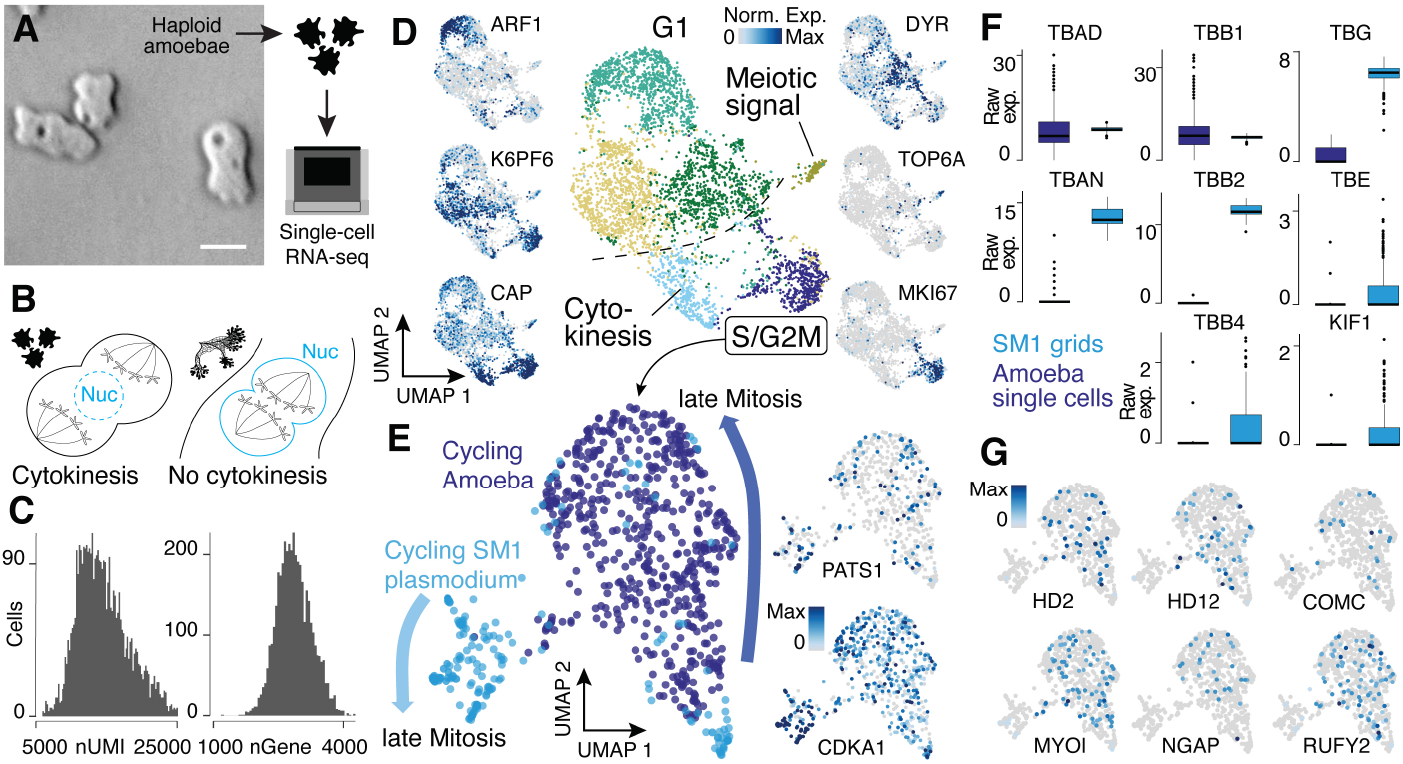
Single-nucleated amoebae are heterogeneous and differentially regulate cytokinetic programs compared with syncytial nuclei. (**A**) Brightfield image of amoebae (Single frame from Supp Movie S3). Scale bar: 10 μm. (**B**) Schematic illustrating mechanistic differences during mitosis in amoebae (left) and plasmodia (right). (**C**) Histograms reveal the distribution of transcript and gene detection counts for amoeba scRNA-seq data. (**D**) UMAP embedding of single haploid amoeba cells (center). Cell cycle stages are indicated and gene expression intensities are visualized as features on the embedding (left/right). (**E**) UMAP embedding of single amoeba cells in G2M phase and samples of a cycling plasmodium (SM1). RNA-seq samples were integrated using Seurat’s built in data ‘anchoring’ (38) to allow a comparison between single-cell (10x Genomics) and bulk RNA-seq (SmartSeq2) data. Feature plots of PATS1 and CDKA1 reveal cell/nuclei cycle directionality for samples. Drawn arrows reveal directionality. (**F**) Cell cycle related marker gene expression identified through differential gene expression analysis is visualized as a boxplot for plasmodium and amoeba samples, respectively. (**G**) Amoeba specific genes are visualized as feature plots on the UMAP embedding in (E).

An interesting feature of the multinucleated syncytia is that the mitotic spindle forms within the nucleus to segregate chromosomes into two daughter nuclei, which is common among syncytial fungi (“closed mitosis”, Fig. 4B) (39–41). In contrast, *Physarum polycephalum* amoebae anchor the mitotic spindle through centrosomes at the opposing poles of cells (“open/astral mitosis”) as occurs during typical eukaryotic cell divisions (Fig. 4B) (39, 42). We therefore integrated spatial transcriptomes from the cycling syncytium (SM1) and transcriptomes from amoeba mitotic cells, and explored the expression similarities and differences. We observed genes that increase expression towards late mitotic phases in both *Physarum* states (Fig. 4E), for example CDKA1 and PATS1 which are involved in cytoskeleton dynamics during cytokinesis (43, 44). However, we observed striking differences in other gene expression profiles. For example, certain tubulins and one kinesin - major components of the spindle assembly pathway, are differentially expressed between the plasmodium and amoebae (Fig. 4F). We find that alpha tubulin 1a (TBAD) and beta tubulin 1 (TBB1) are spindle markers expressed in both states, whereas alpha tubulin 1b (TBAN) and beta tubulin 2 and 4 (TBB2, TBB4) are only detected at high levels in the plasmodium, consistent with previous reports (45, 46). Also, the epsilon and gamma tubulin chains (TBE, TBG) involved in centrosome duplication and spindle formation and the spindle-associated kinesin KIF1, are detected at much higher levels in the plasmodium. Other spindle components such as dyneins do not show a state-specific expression. These data suggest that differential regulation of tubulins orchestrates a change from an open mitosis with cytokinesis in amoeba, to a closed mitosis with nuclei division in the plasmodia, resulting in a syncytia. We note that many amoeba-specific genes could not be attributed to cell cycle pathways. Instead, we observe differential expression of genes related to the amoeba lifestyle (Fig. 4G) such as MYLI involved in phagocytosis and filopodia assembly (47, 48), COMC a gene important for intercellular communication (49), the RAS GTPase activating gene NGAP which is involved in chemotaxis in Dictyostelium (50) and multiple transcription factor-like genes (HD2, HD12, RUFY2).

Together, these data reveal similarities and differences in transcriptome state regulation in amoebal and plasmodial phases of the *Physarum* life cycle.

## Discussion

Multicellular organisms achieve a huge complexity of forms and functions through the compartmentalization and specialization of different parts of the organism. However, the slime mold *Physarum polycephalum* is an extraordinary example of an ostensibly unicellular organism that exhibits specialized morphologies and behaviors across its life cycle. The *Physarum* genome encodes instructions to generate and maintain both single-nucleated amoeboid cells as well as the gigantic single syncytial cell that develops complex structures and exhibits intricate behaviors. Indeed, polycephalum can be translated as the “many-headed” slime, which describes the remarkable ability of *Physarum* to develop multiple foraging growth fronts to sample the environment and seek out new terrain. This structural and behavioral complexity raises the questions around how such specialization and coordination can be generated, maintained and quickly adapted to changing environmental conditions?

Our results suggest that *Physarum*’s dynamic structure formation and response to local environmental conditions is coordinated in part through localized gene regulation. We observed striking spatial differences in gene expression profiles within individual syncytial cells. These patterns appeared to distinguish different structural components of the plasmodium, and also correlated with environmental stimuli (e.g. nutrient source) at individual growth fronts. Our view is that genomic DNA partitioned into nuclei are distributed throughout the syncytium, and serve as local processing units that receive local and systemic input signals to coordinate encoded outputs. Nuclei can be locally fixed in the ectoplasm and remain in the same position for extended time periods, or they can be advected over short or long distances through the endoplasmic shuttle flow as the plasmodium pulsates and grows. This system enables the syncytium to establish complex behavioural responses that can vary in distant parts of the slime mold.

Intriguingly, we find that there is transcriptome heterogeneity among nuclei within the same structure or region. Single-nuclei analysis revealed different proportions of nuclear transcriptome states in the growth front and fan regions, suggesting that it could be beneficial to maintain state variation for rapid access to different transcriptional programs. These data suggest that one mechanism for rapid and diverse response might occur through the co-maintenance of multiple specialized or randomized epigenetic states. We currently do not believe that the syncytium is an assembly of intrinsically specialized nuclei that exist prior to plasmodium formations, as we do not find clear evidence of linked specialized states within *Physarum* amoebae. However, a future area of research will be to understand if transcriptome states are propagated through nuclei divisions such that outputs can be maintained rapidly in the local growth front or network microenvironment. In addition, consistent with previous reports (14), we observed many intact nuclei within the endoplasm. These nuclei shuttle at rapid speeds throughout the plasmodium within a few contraction periods (25, 51) and it is unclear whether such a system could be used to “seed” nuclei states into particular locations within the mold. Interestingly, nuclei in fan regions are 4 times less likely to be dispersed than nuclei in network regions (52), suggesting that nuclei states within fan regions are more likely to be only locally propagated. In addition to local gene regulation, there may also be mechanisms that transport mRNA to particular positions for localized translation as occurs in other single cells with network structures or multiple environmental interfaces (e.g. neurons (53)). However, our data strongly support nuclei adaptation to local and temporally variable environmental stimuli.

It has been hypothesized that the last common ancestor of protists, fungi, and animals was a facultative multicellular organism based in part on shared epithelial structures and cell adhesion catenins being found in all phyla (54, 55). This may suggest that *Physarum* could have lost some ability to establish multicellular structures. Our comparative analysis of single-nucelated amoeba and syncytial cell states reveal that mechanisms for different types of nuclei division are encoded in the *Physarum* genome. Our data also supports that the evolutionary driver for multicellularity is not fully explained by specialization into different cell types, as the *Physarum* plasmodial state reveals that local specialization can be achieved without cellular compartmentalization. The syncytial strategy as is observed in the plasmodial state may be advantageous in certain scenarios, even for cell types within multi-cellular organisms including humans (56). Endoreplication supports rapid cellular and organismal growth and studies in plants suggest that multinucleation and the syncytial strategy may enhance plasticity in response to environmental stress (57). Our data adds to this view by revealing that individual nuclei can be mobilized to process local environmental signals under variable environmental conditions, and that maintaining a pool of nuclei states at all locations could provide rapid and stochastic access to genetic programs at any location within the syncytium.

We also observe that many proteins expressed during syncytial endoreplication have similarities to viral proteins (Fig. S4F-G). This is interesting as it is known that certain viruses such as HIV and SARS-CoV2 can induce syncytial formation (58, 59), and the proteins Syncytin-1/2 in syncytium formation during evolution of the placenta were likely derived from retrovirus infections in mammalian ancestors. In the future, it will be interesting to understand viral contributions to plasmodial formation and evolution in slime molds.

Altogether, our work reveals that slime molds such as *Physarum* offer the extraordinary opportunity to explore how organisms can evolve regulatory mechanisms to divide labor, specialize, balance competition with cooperation, and perform other foundational principles that govern the logic of life.

## Acknowledgements

We thank all lab members of the Treutlein, Alim and Camp labs for helpful discussions and feedback. In particular, we thank Damian Wollny for comments on the RNA extraction procedure. We are grateful for the help of Noah Ziethen and Björn Kscheschinski with sample collection and for the advice of Christian Westendorf on microscopy. We thank Antje Weihmann and Barbara Schellbach for performing sequencing runs and the Single Cell Facility (SCF) at D-BSSE for insights in dyes and imaging approaches.

## Author contributions

TG and CL performed spatial RNA-seq and sc/snRNA-seq experiments and analyzed the data. NS, CL, AJ and SC acquired imaging data with NS analyzing the data with help of AJ. MS and AWei performed FACS. AWeb cultured and prepared slime molds. TG, CL, NS, KA, BT and JGC designed the study, interpreted data and wrote the manuscript.

## Materials and Methods

### Growth and culture of *P. polycephalum*

*Physarum polycephalum* was cultured as microplasmodia in 100 ml liquid culture of 50% semi-defined medium (SDM) and 50% balanced salt solution (BSS), together with hemin, streptomycin and penicillin, while shaking at 180 rpm in a 25°C incubator (60). Change of medium was carried out three times a week. Axenic *Physarum* amoeba culture was grown in 50 ml liquid shaking culture of semi-defined medium (SDM) at 150 rpm and 30 °C. Subculturing happens three times a week to reach a seeding density of 0.2×10^6^ cells/ml (61).

### Sample preparation for fluorescence imaging

For fluorescence imaging, a less auto-fluorescent substrate, 1.2% w/v of non-nutritious PhytagelTM was used and distributed in Petri dishes (Ø = 100mm). Small amounts of microplasmodia (1 ml) were inoculated onto Petri dishes and incubated over 16-24 h in order to grow small 0.5-1.0 cm sized plasmodial networks. For live imaging, nuclei staining was performed by immersing the specimen with 5 μM SYTOTM 62 (Thermo Fisher S11344) dye for 30 minutes and washing with BSS solution. The excitation and emission spectrum (649/680) of this dye is far from the auto-fluorescence spectrum (ex/em 300-450/400-550). To test the performance of SYTO(™) dyes a colocalization experiment on fixed samples was carried out (Fig. S1A-B). Slime molds were first grown with the described procedure. For fixation, 2 ml of 4% PFA solution was added on top of the plasmodium with an incubation time of 15 min. The molds were washed 2 times with autoclaved BSS solution and immersed in a solution of 2.5 μM Hoechst 33342 (Thermo Fisher 62249) and 0.5 μM SYTOX Deep Red (Thermo Fisher S11380) for 30 minutes. The ex/em ranges of the dyes are 361/497 and 660/682, respectively. To map the nuclei density of whole plasmodia, *Physarum* was grown from microplasmodia between a thin Phytagel(™) sheet and the Petri dish bottom. This is to limit and homogenize the height of the plasmodia for better comparison of nuclei densities. A blade was used to run along the perimeter of the Phytagel(™)-overlay to release the thin sheet from the bottom, before placing the microplasmodia under it. To ensure the immersion of fixation solution and dye will reach the specimen, the staining-protocol was also adjusted. PFA solution (4%, 2 ml) was placed on top of the Phytagel(™) and the petri dish was shaked it on a tilting bed (Polymax 1040, Heidolph Instruments) at 2rpm for 45 minutes, before washing it 3 times with BSS solution. For fluorescent labelling approximately 200 μl of 2.5 μM Hoechst33342 were used. The stained specimen was first placed on the tilting bed for 30 minutes at room temperature and subsequently stored in the incubator at 25°C overnight. Finally, the slime mold was washed twice with BSS.

#### DAPI sample preparation

Slime molds were grown on regular agar plates as described above. 4% of PFA solution was added on top of the plasmodium with an incubation time of 30 min. The molds were washed 2 times with autoclaved distilled water and DAPI (Thermo Fisher D1306) (300 nM) was added on top. The molds were incubated for 10 min with the staining solution. The solution was then removed and the molds were washed 2-3 times with autoclaved distilled water.

### Fluorescence imaging

Microscopy images were acquired with the Zeiss Axio Zoom V16 stereo microscope with a Zeiss Plan Neofluar Z 2.3x/0.57 objective, a HXP 120C fluorescence lamp and a Hamamatsu Orca Flash 4.0 V2 complementary metal-oxide semiconductor (CMOS) camera. We use ZEN Blue Edition (Zeiss) to control the microscope. To image Hoechst and red SYTO dyes, filter cubes 02 (BP 365/ FT 395/ LP 420) and 50 (BP 640/ FT 660/ BP 690) were used, respectively. For reference autofluorescence imaging filter cube 38HE (BP 470/ FT 495 / BF 525) was used. To improve z-resolution we use the Zeiss ApoTome. 3D micrographs of whole living and fixed cells were acquired directly after staining, by scanning over the organism using a motorised xy stage. For every tile of the micrograph a bottom-to-top z scan was performed over the fluorescent and the autofluorescent channels, with a z-step of 2 - 5 μm and a magnification of 65-258x.

#### DAPI imaging

Images were acquired on a Leica TCS SP5 with the following settings: laser power at 29.08, 10.0x × 0.30 DRY HC PL FLUOTAR objective with a numerical aperture (NA) of 0.3 and UV lenz FW 20x/ 0.70. Images were generated directly after DAPI staining and were acquired from the bottom through microscopy slide and agar layer. DAPI was excited at 405 nm and was measured with the filter set Leica/DAPI 430/550. The scan speed was at 200 ms, the pinhole at 7.07×10-5 and the pinhole airy at 0.99. The zoom-in was performed by increasing the dbl zoom property from 1 to 3 at plasmodial regions of interest.

### Image Processing

A custom written Fiji macro is used to deconvolve the z stack of the SYTO channel for each tile of the micrograph using the Fiji plugin deconvolutionlab2 (62) and a generated point-spread-function using PSF-Generator (63). After deconvolving the z-stacks we maximum-intensity project the images and use a convolutional neural-network based StarDist-model (18) to predict the position and shape of nuclei within the micrographs. Accuracy of the model was tested using a ground-truth dataset training of a Weka-classifier (64) with subsequent curation of the segmentation. We find an overall precision (TP/(TP+FP)) of 94% and accuracy of 81% (TP/(TP+FP+FN)) indicating a slight underestimation of particle number. Using the Microscopy Image Stitching Tool (MIST) (65) we stitch the tiles of the auto-fluorescence micrograph together, which serves as a reference stitching scheme for the fluorescence micrographs and the segmented images. The resulting label-image of the fluorescent spots is analysed for the object area, eccentricity and diameter. Furthermore, we use the detected spot positions to analyze internuclear distance using a Ball-Tree algorithm and to perform a Kernel Density Estimation (KDE). The resulting probability density function is integrated over small regions to measure the nuclei-density within the slime mold (Fig. S1C).

### Growth and preparation of the plasmodium for sequencing

The slime molds used for sequencing were all provided by the lab of Karen Alim at the Max Planck Institute for Dynamics and Self-Organization (Göttingen, Germany). All slime mold plasmodia originated from microplasmodia liquid cultures as described recently (60). Briefly, plates had a small cavity in the agar where a small volume of microplasmodia culture was dispensed. In most of the cases, the microsplasmodia needed around 1 day to generate a network structure and expanding as plasmodia outside of the cavity. Oat flakes were added to the plates one or two days after if the experimental design required it.

### Preparation of nuclei and cell suspensions for 10x Genomics experiments

Samples were kept on ice during all steps of the nuclei preparation. Slime molds were incubated in 2 ml homogenisation buffer (10 mM CaCl2, 0.1% Nonidet P-40, 10 mM Tris-HCl, pH 8, 0.25 M Sucrose) for 5 min. In addition, the suspension was pipetted up and down a few times to help dissociating the tissue. Afterwards, samples were transferred to a douncer and 15 strokes with pestle A and another 15 strokes using pestle B were applied. A 100 μm strainer was used to filter the suspension and was afterwards flushed with 2 mL HBSS to collect remaining nuclei. Nuclei were spun at 300 g for 5 min. After removing the supernatant nuclei were resuspended in 500 μl HBSS and filtered through a 20 μm strainer. Another 500 μl HBSS were used to wash off remaining nuclei from the filter membrane. DAPI (1:1000) was added to the nuclei suspension and FACS sorting (85 μm nozzle) was performed to isolate single nuclei. 85,000 single nuclei were collected in a well of a 96 cell-culture plate and the concentration of the nuclei, ranging between 200 and 300 nuclei/μl, was estimated using a hemocytometer. FACS sorting of nuclei extracted from amoeba was not possible due to the sensitivity of the nuclei and the single-cell suspension of amoeba was therefore directly used for 10X Genomics experiments. Cells or nuclei were loaded onto the 10X cartridge (Chip B) at various concentrations, with loading the maximum volume possible for nuclei suspensions or targeting 6000 cells for the amoeba suspensions. Cell encapsulation, cDNA generation, and preamplification as well as library preparation were performed by using the Chromium Single Cell 3’ v3 reagent kit according to the kit protocol.

### Spatial transcriptomics of plasmodia

100 μL of microplasmodia were plated on a lid of a Corning® 384-well Clear Bottom Polystyrene Microplates after spreading a thin layer of agar across the inner side of the lid. The inner side of the lid was covered with parafilm and incubated at room temperature in the dark for 20 hours. On the next day the 384-well plate was prepared by adding 20 μl lysis solution (6 M Guanidine Hydrochloride, 1% Triton X-100) in each well (adapted from (66)). Lids with plasmodia spanning a large area of the lid were placed on the 384-well plate and samples were collected by spinning the plate at 4000 g for 2 minutes. After sample collection the plate was placed on dry ice until further processed or frozen at −80 degrees. Note that the collection of slime mold pieces for the initial test experiments (Fig. S3) happened manually by sampling individual 2×2 mm grids of 3 different plasmodia one after another (Fig. S3B). A 96-well PCR plate (Eppendorf) was prepared by adding 13 μL of Agencourt RNA XP SPRI beads to as many wells as needed. RNA was extracted from the lysis mix by transferring 5 μL of lysis mix from the 384-well plate to the 96-well plate prepared with RNA XP SPRI beads (bead to sample ratio of 2.6x; vol:vol). The suspension was incubated for 5 minutes at room temperature. After binding beads at the tube walls using a magnetic rack the supernatant was removed. After two rounds of washing with 80% ethanol, RNA was released from the beads by resuspending them in 10 μl EB buffer. Drying the beads was not needed and the purified RNA suspension was transferred to a fresh 96-well plate and stored at −80 degrees until further processed. 2 μl of this bulk RNA was used as input for SmartSeq2 (20). Generation of cDNA followed SmartSeq2 instructions with 16x PCR cycles for cDNA amplification. A high-throughput electrophoresis-based fragment analyzer (Fragment Analyzer, Advanced Analytical Technologies) was used to assess the cDNA fragment size distribution and its concentration of exemplary samples. Illumina libraries were constructed by using the Illumina Nextera XT DNA sample preparation kit according to the protocol. Up to 192 libraries were pooled (3 μL each) and purified with Agencourt SPRI select beads.

### Sequencing

Library concentration and size distribution were assessed on an Agilent Bioanalyzer and with Qubit double-stranded DNA high-sensitivity assay kits and a Qubit 2.0 fluorometer for all sequencing libraries. 10xGenomics sequencing libraries were sequenced following the 10xGenomics protocol to a depth of 50,000 to 100,000 reads per nucleus or cell. Base calling and demultiplexing of single nuclei or cells were performed by using 10xGenomics Cell Ranger 3.1 software. SmartSeq2 libraries were paired-end sequenced (100 base reads) on an Illumina HiSeq 2500 aiming for a depth of 200,000 reads per sample, base calling was performed using Bustard (67) and adaptor trimming and de-multiplexing were performed as described previously (68).

### Data Analysis

All sequenced datasets were aligned to the latest version of *Physarum*’s transcriptome (34). All raw reads were aligned using STAR (69). Mapping of reads of the 10x Genomics data sets was performed using Cell Ranger 3.1 implemented in the 10x Genomics analysis software, generating absolute transcript counts based on unique molecular identifiers (UMI). Reads obtained for the SmartSeq2 data sets were mapped with a local installation of STAR and TPM values were estimated using a custom Perl script. Customised R Studio scripts (see https://rstudio.com) were used for final preprocessing steps and to run all data analyses. Scripts can be obtained upon request. The R packages ‘Seurat’ v3 (38) and ‘ggplot’ were used for almost all analyses and visualizations unless stated differently. For all feature plots created by Seurat, feature-specific contrast levels based on quantiles of non-zero expression were calculated (q10/q90 cutoff).

#### Spatially resolved bulk RNA-seq analysis

SmartSeq2 RNA-Seq grid data sets were first merged per plasmodium and log2 TPM values were calculated by transforming all values with TPM > 1 and setting all values with TPM < 1 to 0. Grids with small read coverage (Paired reads SM1<200k, SM2-4<100k) or no clear overlap with the underlying plasmodium network were removed. Only transcripts with a UniProt annotation or at least a gene description in the UniProt annotation file by Glöckner and Marwan (34) were kept for further analyses while removing all other genes from the grid/gene matrices. Each data set was then loaded and processed separately using Seurat (Seurat’s default normalization was omitted) and data scaling was performed with nGene and nRead regression. Dimensionality reduction consisted of a principal components analysis (PCA) and Uniform Manifold Approximation and Projection (UMAP) with the first 5-15 (SM1:15, SM2:5, SM3:5, SM4:15, T1:10, T2:10, T3:5) principal components (PCs). Louvain clustering (Seurat default) was performed for the same number of input PCs with a resolution of 0.1-0.6 (SM1:0.4, SM2:0.1, SM3:0.3, SM4:0.2, allT:0.6). For plasmodium SM4 and T1 Louvain clustering gave no convincing results and we therefore hierarchical clustered grids using hclust’s ward method with distances estimated by Pearson correlation and the same input gene sets as described above. Differential gene expression analysis was performed using the Wilcox test (Seurat default) on identified clusters or on a selected region for SM4 (Fig. 2H). The DE genes identified for the oat flake covered grids were visualized as Volcano plot using the R package ‘EnhancedVolcano’ (https://github.com/kevinblighe/EnhancedVolcano). For the density plots of grid distance against transcriptome correlation we selected top 200 variable genes per individual plasmodium and extracted the scaled matrix (genes/grids) using these genes from each plasmodium. Scaled values were converted into purely positive values by adding the absolute minimum of a gene to the scaled values across all grids for a given gene, respectively. The obtained matrix was then correlated against itself with ignoring diagonal grid correlations. Distances between grids were calculated within the cartesian scatter by using the R package ‘raster’. The density plots were generated using R’s smoothScatter() function by plotting grid’s densities against transcriptome correlations. The regression line was calculated as a linear model. Seurat objects were afterwards merged to obtain a combined data set and dimensionality reduction was performed as described above with 5 input PCs for the UMAP embedding and Louvain clustering with a resolution of 0.2.

#### Analysis of snRNA-seq plasmodium data

Only transcripts with a UniProt annotation or at least a gene description in the UniProt annotation file by Glöckner and Marwan (34) were kept for further analyses while removing all other genes from the nuclei/gene matrix. Afterwards, single-nuclei transcriptome information was loaded into R Studio using Seurat. Datasets corresponding to the primary plasmodium were merged and nuclei with less than 3000 detected counts were used for further analyses using the Seurat package. The same process was accomplished for the datasets corresponding to the secondary plasmodium. The datasets were log normalized using the Seurat function (scale factor of 10,000). Neither feature selection nor mitochondrial gene removal were performed and the data was directly standardised, by regressing out differences in UMI and gene detection counts (Seurat package). Technical variation between runs within the same experiment was identified to be neglectable. The dimensionality reduction process consisted of a principal components analysis (PCA) and Uniform Manifold Approximation and Projection (UMAP) with the first 15 principal components (PCs). Louvain clustering (Seurat default) was performed for the top 15 PCs with a resolution of 0.3. For the primary plasmodium we identified cluster 3 as purely driven by low nUMI and nGene counts and we therefore removed this cluster from the data set. Afterwards we re-clustered and re-embedded the data set using only top 7 PCs and a resolution of 0.4. All steps mentioned above were performed using the functions from the Seurat package. Differential gene expression analysis was performed using the Wilcox test (Seurat default) on identified clusters. Transcripts and cluster markers were queried to the UniProt annotation file obtained from Glöckner and Marwan (34). Cycling nuclei were extracted by cluster (1 and 2) and only marker genes identified for the 3 clusters of the spatial transcriptomic data set of SM1 were used as input for PCA. Afterwards the top 5 PC components were used for UMAP embedding and Louvain clustering. Pseudotemporal ordering of the cycling nuclei was performed using a diffusion map algorithm of the R package destiny (70). Raw ordering results were separately obtained depending on the UMAP location ([c4],[c0/c2/c3],[c1]) and afterwards sorted and ranked. Gene expression is visualized as a smoothed LOESS function (span = 10) against the pseudotemporal ordering. Pseudobulk estimates were obtained for each cluster by averaging each gene’s normalized expression value across nuclei of the respective clusters. The same was done for the grid based RNA-seq clusters of SM4. The pseudobulk matrix of the nuclei data was afterwards correlated against the pseudobulk matrix of the grid based spatial data using Spearman’s correlation including only top 50 cluster marker genes identified in the spatial RNA-seq data set of SM4. Correlation values were scaled first by column and then by row and values were cut to 1 or −1 when being either larger or smaller, respectively. Correlation values were used to hierarchical cluster the nuclei clusters using the ward method after calculating distances with Pearson correlation. Afterwards we selected two clusters (c0 and c4) and correlated the nuclei pseudobulk data against individual grids of SM4 using Spearman correlation and the same gene set as used above. Raw correlation values were visualized as features on the grids and can be interpreted as either fan or network scores.

#### Analysis of scRNA-seq amoeba data

Again, only transcripts with a UniProt annotation or at least a gene description in the UniProt annotation file by Glöckner and Marwan (34) were kept for further analyses while removing all other genes from the cell/gene matrices. Datasets of the 2 sequencing lanes were merged using Seurat and cells with detected UMI counts ranging from 6,000 to 25,000 were extracted for downstream analyses. Normalization and standardization steps were performed identically as described above. Neither feature selection nor mitochondrial gene removal were done. PCA was followed by UMAP and Louvain clustering with the first 25 PCs and a resolution of 0.6. Differential gene expression analysis was performed using the Wilcox test (Seurat default) on identified clusters. Cycling amoebae were extracted by cluster identity (cluster 3) and afterwards merged with the Seurat object containing the spatially resolved transcriptomes of SM1. Data integration was performed using Seurat’s built in function for ‘anchoring’ data of different experiments to remove the effect of single-cell platforms used during data generation (38). We used all genes for finding anchors between the data sets and afterwards 20 PCs for the integration with the identified anchors. The UMAP embedding and Louvain clustering was then performed on the integrated data using top 10 PCs and with a resolution of 0.4 for clustering.

---

**Fig. S1.**
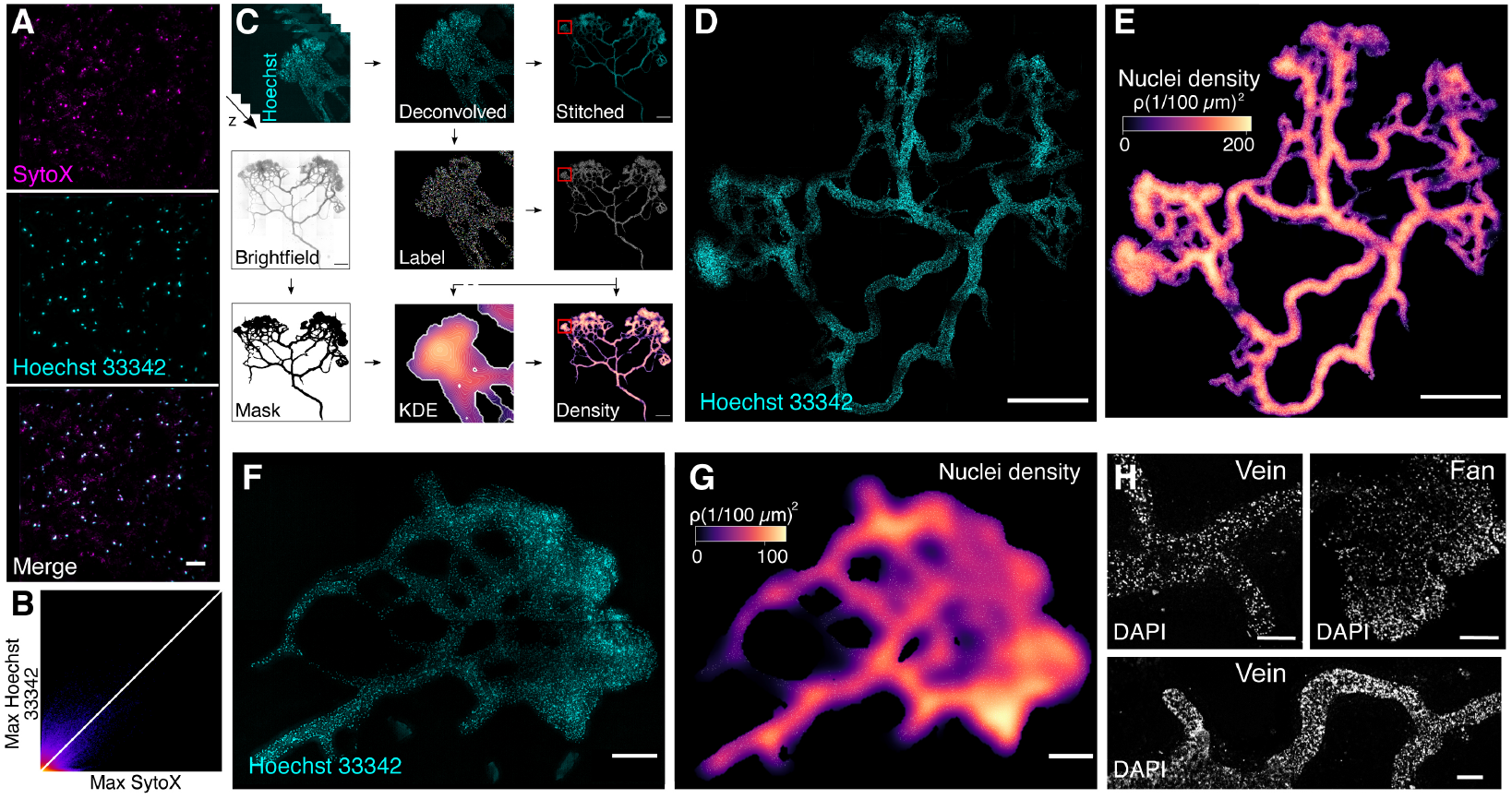
Nuclei are not homogeneously distributed across the multinucleated plasmodium. (**A**) Colocalization tests of Hoechst and Syto. Maximum projections are shown for SytoX, Hoechst 33342 and the merged image (top to bottom). Scale bar: 50 μm. (**B**) Maximum 2D projection histogram revealing high correlation of Syto and Hoechst (Pearson’s R value 0.73). White line indicates R=1. (**C**) Methodology workflow for imaging data analysis. (**D**) Hoechst stained nuclei across a fixed plasmodium. Scale bar: 1000 μm. (**E**) Density map of nuclei for the plasmodium shown in B after StarDist segmentation and kernel density estimation. Scale bar: 1000 μm. (**F**) Hoechst stained nuclei across a fixed plasmodium. Scale bar: 200 μm. (**G**) Density map of nuclei for the plasmodium shown in D after StarDist segmentation and kernel density estimation. Scale bar: 200 μm. (**H**) DAPI stained nuclei across a fixed plasmodium. Different regions of the plasmodium are shown. Scale bar: 500 μm.

**Fig. S2.**
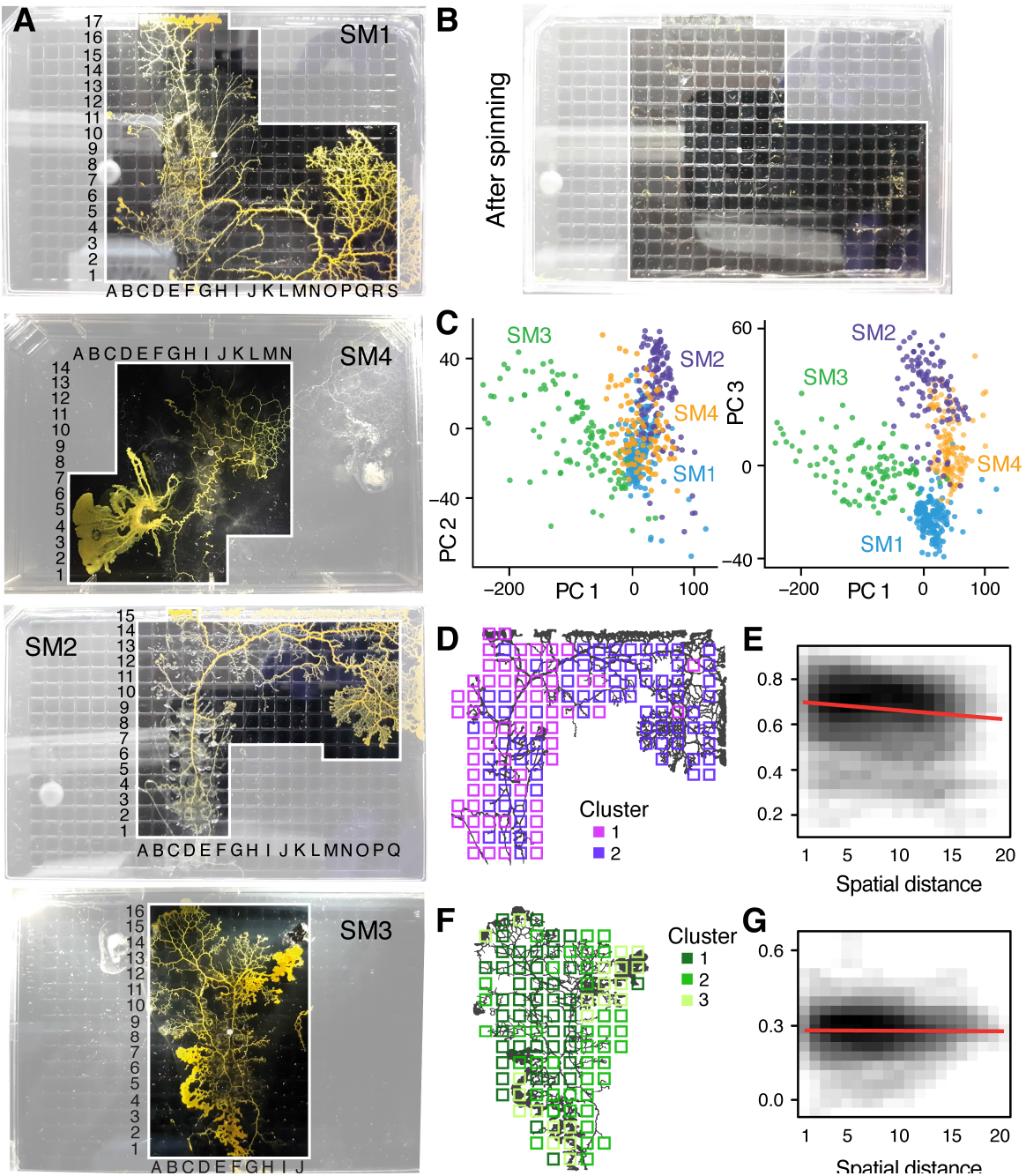
Spatial Transcriptomics experiments allow to identify gene expression differences across plasmodial slime mold syncytia. (**A**) Camera images showing 4 slime mold plasmodia just before spun into 384-well plates, respectively. Oat flakes had been removed for SM3 and 4. (**B**) Exemplary image of a 384-well plate after spinning SM1 into separate wells of the plate. (**C**) PCA results for the combined RNA-seq data set of all 4 slime molds. Each dot represents a single sample (well of the 384 well plate). (**D**) Sampling grid on the 384-well plate overlaid by the plasmodium of SM2 at the time when sampled. Different colors represent the clustering result from (Fig. 3D). (**E**) Density plot visualizes the relation between the pairwise distance against the pairwise Pearson correlation across all grids of SM2. A statistically significant negative correlation is identified and marked by a red line. (**F**) Sampling grid on the 384-well plate overlaid by the plasmodium of SM3 at the time when sampled. Different colors represent the clustering result from Fig. 3D. (**G**) Density plot visualizes the relation between the pairwise distance against the pairwise Pearson correlation across all grids of SM3. No clear correlation is identified as indicated by a red line.

**Fig. S3.**
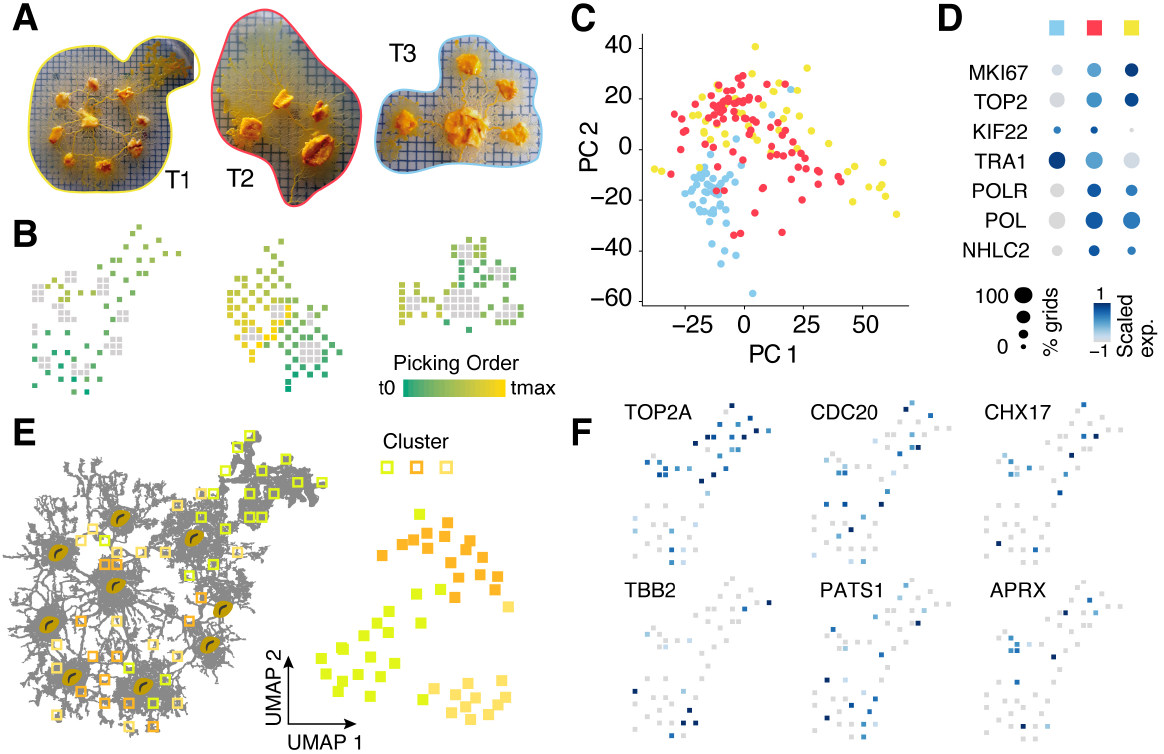
Spatial Transcriptomics experiments with manual picking of plasmodial pieces. (**A**) Camera images showing 3 slime mold plasmodia prior to consecutively picking individual pieces of roughly 2×2 mm into separate reaction tubes while keeping track of the picking location. (**B**) Sampling grids across the plasmodia in (A) with oat flakes visualized as grey squares to allow an optical overlay with the images in (A). The picking order is visualized as a feature onto the grids. (**C**) PCA scatter plot of PC1 and 2 reveals separation of plasmodium T3 from the other 2 plasmodia. Same color legend as in A is depicted. (**D**) Dotplot representation of nuclei division genes that are identified to be top marker genes of plasmodium T1 and T2. Same color legend as in A is depicted. (**E**) Left: Sampling grids are visualized as an overlay with the shape of plasmodium T1 and are colored by the clustering result. Right: UMAP embedding of sampling grids of T1. Colors depict cluster identity. (**F**) Gene expression is visualized as feature on the UMAP embedding in (E). Early (TOP2A), medium (CDC20) and late (TBB2/PATS1) nuclei division cycle markers are shown. Coexpression of CHX17 and APRX is found in a region with high oat flake density.

**Fig. S4.**
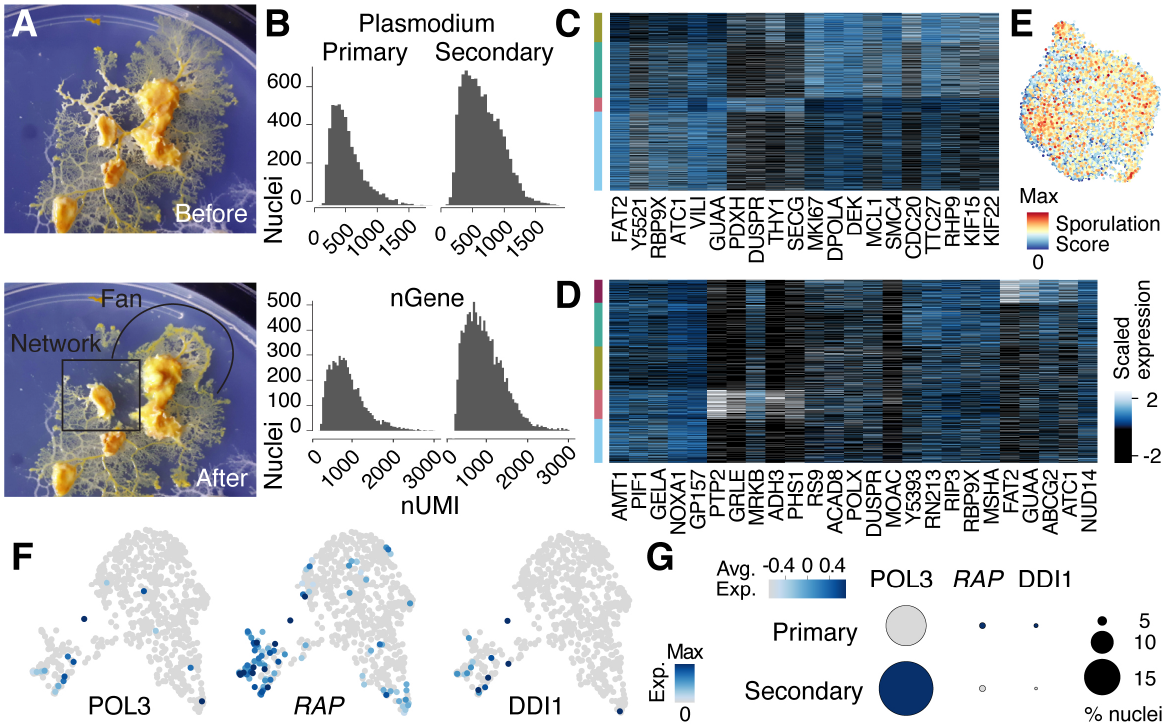
Nuclei heterogeneity revealed by single-nuclei RNA-seq. (**A**) Camera images show the secondary plasmodium before and after sampling a fan and network enriched sample (encircled by black lines), respectively, for the snRNA-seq experiment. (**B**) Histograms visualize transcript and gene detection counts per nucleus for each plasmodium sampled. (**C**) Heatmap representation of marker gene expression for primary plasmodium clusters shown in Fig. 3B. (**D**) Heatmap representation of marker gene expression for secondary plasmodium clusters shown in Fig. 3E. (**E**) Sporulation score is shown as a feature onto the UMAP embedding in Fig. 3E. The score was estimated as the sum of transcript counts for a list of fruiting body formation specific genes (34). (**F**) Gene expression feature plots on UMAP embedding of Fig. 4E (Integrated SM1 grids and single-cell amoeba) revealing plasmodial specific expression of virus-related genes (RAP = Retroviral aspartyl protease domain). (**G**) Dotplot visualization of virus-related genes in (F) for primary and secondary plasmodium single-nuclei RNA-seq data revealing expression also in sparse nuclei data with 2 of them being more highly expressed in the cycling primary plasmodium.

